# *Butyrivibrio hungatei* MB2003 competes effectively for soluble sugars released by *Butyrivibrio proteoclasticus* B316^T^ from growth on xylan or pectin

**DOI:** 10.1101/395103

**Authors:** Nikola Palevich, William J. Kelly, Siva Ganesh, Jasna Rakonjac, Graeme T. Attwood

## Abstract

Rumen bacterial species belonging to the genera *Butyrivibrio* are important degraders of plant polysaccharides, particularly hemicelluloses (arabinoxylans) and pectin. Currently, four distinct species are recognized which have very similar substrate utilization profiles, but little is known about how these microorganisms are able to co-exist in the rumen. To investigate this question, *Butyrivibrio hungatei* (MB2003) and *Butyrivibrio proteoclasticus* (B316^T^) were grown alone or in co-culture on the insoluble substrates, xylan or pectin, and their growth, release of sugars, fermentation end products and transcriptomes were examined. In single cultures, B316^T^ was able to degrade and grow well on xylan and pectin, while *B. hungatei* MB2003 was unable to utilize either of these insoluble substrates to support significant growth. Co-cultures of B316^T^ grown with MB2003 revealed that MB2003 showed almost equivalent growth to B316^T^ when either xylan or pectin were supplied as substrates. The effect of co-culture on the transcriptomes of B316^T^ and MB2003 was very marked; B316^T^ transcription was largely unaffected by the presence MB2003, but MB2003 expressed a wide range of genes encoding carbohydrate degradation/metabolism and oligosaccharide transport/assimilation in order to compete with B316^T^ for the released sugars. These results suggest that B316^T^ has a role as an initiator of the primary solubilization of xylan and pectin, while MB2003 competes effectively as a scavenger for the released soluble sugars to enable its growth and maintenance in the rumen.

**IMPORTANCE:** Feeding a global population of nine billion people and climate change are the primary challenges facing agriculture today. Determining the roles of rumen microbes involved in plant polysaccharide breakdown is fundamental to understanding digestion and maximizing productivity of ruminant livestock. *Butyrivibrio* are abundant rumen bacteria and are a substantial source of polysaccharide-degrading enzymes with biotechnological applications for the depolymerization of lignocellulosic material. Our findings suggest that closely related species of *Butyrivibrio* have developed unique strategies for the degradation of plant fibre and the subsequent assimilation of carbohydrates in order to coexist in the competitive rumen environment. The identification of genes related to their enzymatic machinery by which these bacteria work in concert to degrade these different forms of polysaccharides contributes to our understanding of carbon flow in the rumen.

## INTRODUCTION

The bacterial community responsible for plant fibre degradation within the rumen is diverse (1-3), and interactions between bacterial species facilitate this process (4). Bacterial species belonging to the genus *Butyrivibrio* are metabolically versatile (2), and efficiently utilize the insoluble, complex polysaccharides, xylan and pectin (5-8). At present, there are four recognized *Butyrivibrio* species (9), but knowledge of the interactions between these species, and the consequences of these interactions to fibre degradation in the rumen, are poorly understood.

Comparison of the *B. hungatei* MB2003 (10, 11) and *B. proteoclasticus* B316^T^ (8) genomes, and in particular their CAZyme profiles, show that these *Butyrivibrio* species are functionally similar, and predict important roles for these species in the breakdown of hemicelloses and pectin. B316^T^ contains 342 predicted CAZymes, and shows strong growth and rapid degradation of both oatspelt xylan and apple pectin (12). However, phenotypic analysis of MB2003 showed it cannot grow on these insoluble substrates (10, 11), even though it encodes 225 CAZymes associated with hemicellulose and pectin degradation. Therefore, it is hypothesized that MB2003 does not directly hydrolyze xylan or pectin, but rather relies on other rumen organisms with more developed polysaccharide degrading abilities to initiate the degradation of insoluble substrates, and competes for the released oligosaccharides and sugars to enable its growth. To investigate this hypothesis, MB2003 and B316^T^ were grown on oatspelt xylan or apple pectin, separately in mono-cultures to compare their individual substrate utilization abilities, and together in co-cultures, to investigate the interactions that occurred. Growth of the strains was followed with strain-specific qPCR and monosaccharides release from these polymers and their subsequent utilization was measured, along with fermentation end products and gene transcript abundances.

## RESULTS

### *B. hungatei* MB2003 competes with *B. proteoclasticus* B316^T^ for xylan and pectin

*Butyrivibrio hungatei* MB2003 and *Butyrivibrio proteoclasticus* B316^T^ strains were grown as mono- or co-cultures on the insoluble substrates oatspelt xylan and apple pectin, to examine the interaction between the strains. The qPCR analysis of the mono-cultures showed that B316^T^ was able to utilize well, both xylan and pectin for growth (Fig. 1). MB2003 showed only slight growth on pectin, and very little growth on xylan. In competition, co-cultures of MB2003 and B316^T^ displayed similar growth until 16 h when either xylan or pectin were supplied as substrates. B316^T^ grown in co-culture with MB2003 showed greatly reduced growth compared to its growth in mono-culture.

**Figure. 1.**
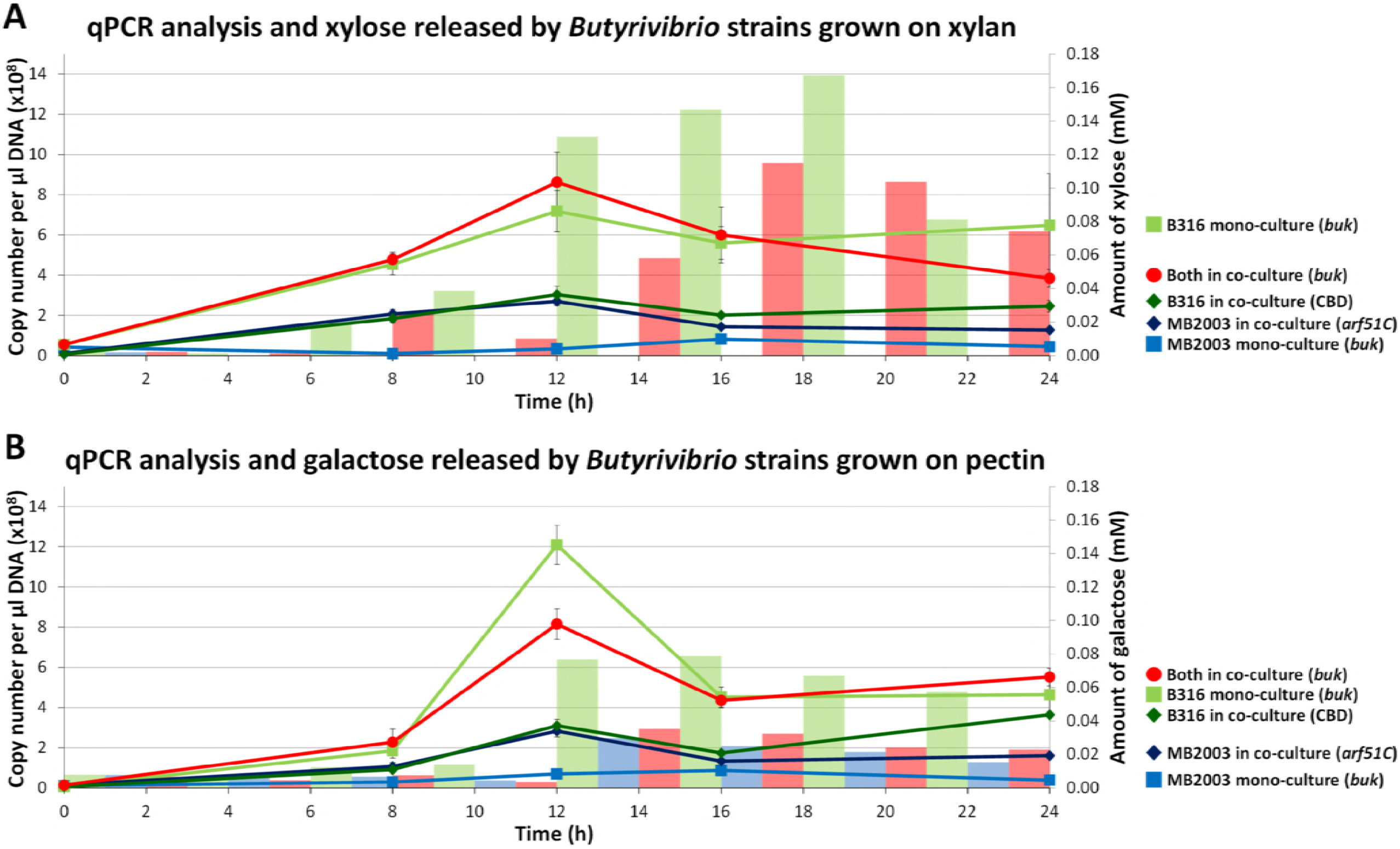
Mono- and co-culture growth of *B. hungatei* MB2003 and *B. proteoclasticus* B316^T^ grown on xylan and pectin as determined by qPCR. A, xylan-grown cultures and xylose released by MB2003 (blue bar) and B316^T^ (green bar) in mono- and co-cultures (red bar). B, pectin-grown cultures and galactose released by MB2003 (blue bar) and B316^T^ (green bar) in mono-and co-cultures (red bar). Error bars represent ±1 standard deviation of the mean.

Fermentation end-products were used as an additional indicator of growth (Table S2). Formate was the main VFA produced in B316^T^ mono- and B316^T^ + MB2003 co-cultures on xylan, followed by butyrate, then acetate. In contrast, acetate was the main VFA produced by B316^T^ mono- and B316^T^ + MB2003 co-cultures grown on pectin, followed by formate, then butyrate. For MB2003 grown on xylan or pectin in mono-culture, only small amounts of VFAs were produced, reflecting the poor growth of this strain alone on either of these insoluble substrates. The total amount of VFAs produced in pectin co-culture samples was less than the combined amounts of VFAs produced by MB2003 and B316^T^ mono-cultures, while co-cultures grown on xylan produced more total VFAs than the mono-cultures combined.

Monosaccharides released from xylan and pectin during growth were measured using ion chromatography (IC; Fig. 1). For B316^T^ cells grown on xylan, the most abundant sugar detected was xylose. Maximum concentrations of xylose were detected at 12 h and 10 h in mono- and co-culture samples respectively, with small amounts of arabinose and galactose detected from 8 to 16 h of growth (Fig. 1A). The xylan-grown MB2003 mono-culture samples showed very little release of monosaccharides (Fig. 1A). The pectin grown cultures showed similar sugar release dynamics in the mono- and co-culture samples (Fig. 1B), but the monosaccharide detected at the highest concentration was galactose, with smaller amounts of arabinose and rhamnose. The B316^T^ mono-culture produced a rapid release of galactose (0.08 mM) from 6 h to 8 h (Fig. 1B), while MB2003 released smaller amount of galactose (0.03 mM) during the same phase of growth (Fig. 1B). A similar amount of galactose (0.04 mM) was released in the co-cultures (Fig. 1B). The results indicate that B316^T^ is an efficient primary degrader of both xylan and pectin, and that MB2003 is able to compete with this bacterium by utilizing small monosaccharides, xylo- and pectic- oligosaccharides.

### Transcriptional changes and differential gene expression

Samples of mono- and co-cultures of B316^T^ and MB2003 grown on xylan or pectin were collected at mid-log growth (12 h) and RNAs were extracted for transcriptome analyses (Data Set S1 and S2). Transcripts from a total of 1,900 genes in B316^T^ and 3,000 genes in MB2003 were detected, and among these 1,300 and 2,206 genes had functional annotations in B316^T^ and MB2003, respectively. The total numbers of differentially expressed genes (DEGs; FDR *Q-value* < 0.05, KW adjusted *p-values* of < 0.05 and ≥ 2-fold log_2_-transformed signal intensity difference; see Text S1) were 17 and 307 for the xylan growth condition, 56 and 787 for the pectin growth condition in B316^T^ and MB2003 respectively (Table S4). The most abundant and DEGs (mono-vs co-culture) for B316^T^ transcripts were those associated with post-translational modifications, carbohydrate metabolism (various CAZymes), sugar ABC transport systems, flagella biosynthesis and some transposases (Data Set S1). In MB2003 the DEGs were associated with acetyl-CoA metabolism, sugar ABC transport system, DNA replication, carbohydrate metabolism (various CAZymes) and flagella biosynthesis (Data Set S1). The full gene annotations, CAZy families, and COG functional assignments of all the DEGs are shown in Data Sets S3 to S6.

During growth on xylan, B316^T^ had 10 genes up-regulated in the mono-culture conditions and 7 genes up-regulated in the co-culture conditions (Data Set S2). In contrast, MB2003 cells grown in mono-cultures on xylan had 215 genes up-regulated while 92 genes were up-regulated in the co-culture condition (Data Set S2). MB2003 also had 14 genes significantly up-regulated in the xylan mono-culture that were not expressed at all in the co-culture situation, and 4 genes in the co-culture conditions that were not expressed at all when grown in mono-culture. During growth on pectin, B316^T^ up-regulated 49 genes in mono-culture while only 7 in co-culture conditions (Data Set S3). There were 26 genes that were only expressed in the mono-culture and there were no genes induced only in the co-culture. MB2003 grown on pectin was very different in terms of DEGs; 725 genes were up-regulated in the mono-cultures and 62 were up-regulated under co-culture growth (Data Set S3). MB2003 had 107 genes up-regulated in the mono-culture that were not expressed at all in the co-culture, and no genes were expressed exclusively in the co-culture. COG classifications were assigned to 56% and 66% of B316^T^ DEGs from cells grown on xylan and pectin, respectively, while 75% and 63% of MB2003 DEGs from cells grown on xylan and pectin, respectively were assigned to COGs. For MB2003 grown on xylan, the number of co-culture DEGs assigned to the carbohydrate metabolism category (G) was similar to the mono-culture, while the number of co-culture DEGs assigned to the cell motility COG category (N) was three times higher than the mono-culture (Fig. 2). This suggests that MB2003 up-regulates its motility functions when co-cultured with B316^T^ on xylan. For the pectin grown cultures of MB2003, DEGs belonging to every COG category analyzed were up-regulated in the mono-culture to a significant extent compared to the co-culture (Fig. 3).

**Figure. 2.**
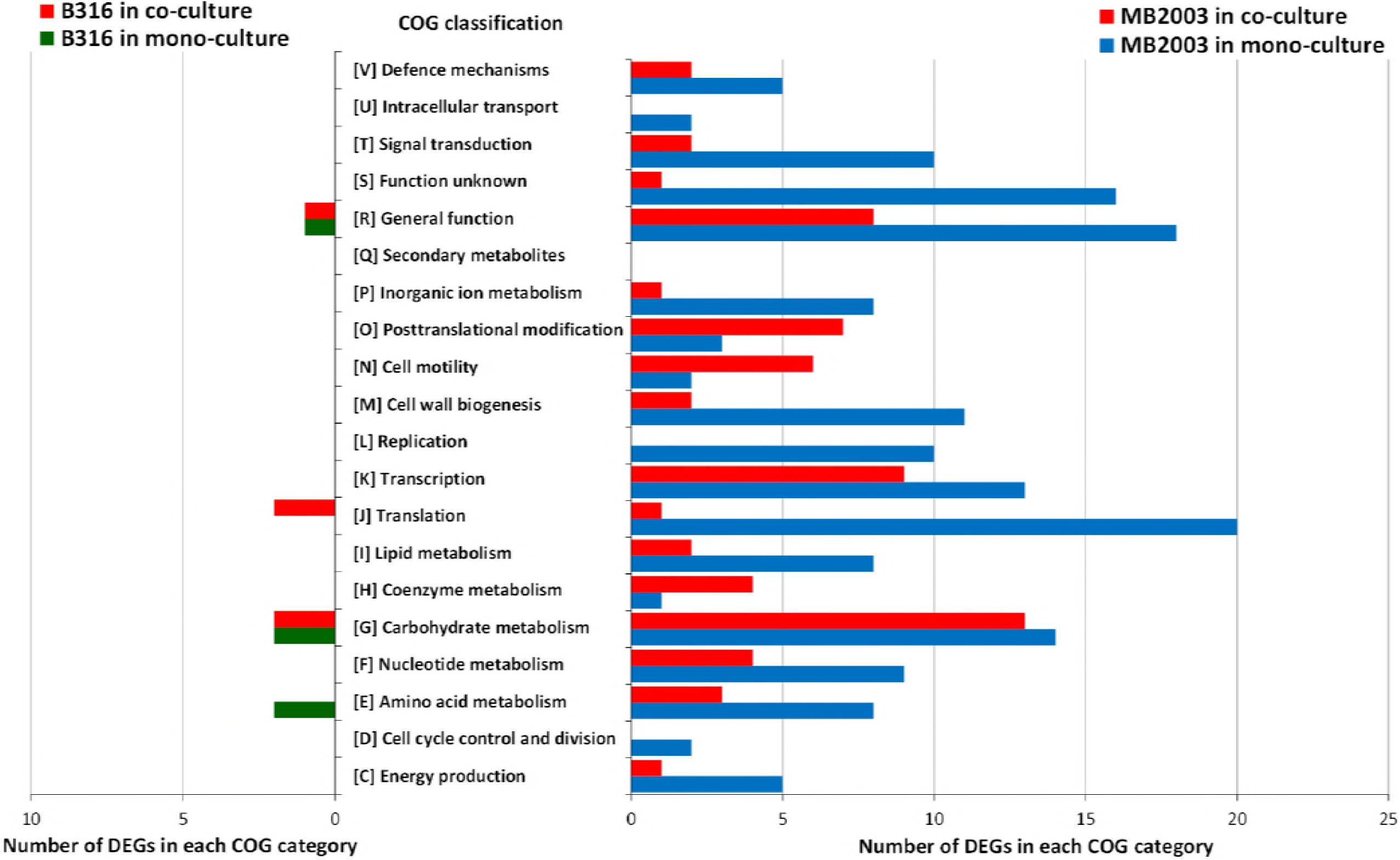
COG classifications of DEGs from *B. proteoclasticus* B316^T^ and *B. hungatei* MB2003 grown in mono- and co-culture on xylan. Analysis includes all DEGs with COG classifications.

**Figure. 3.**
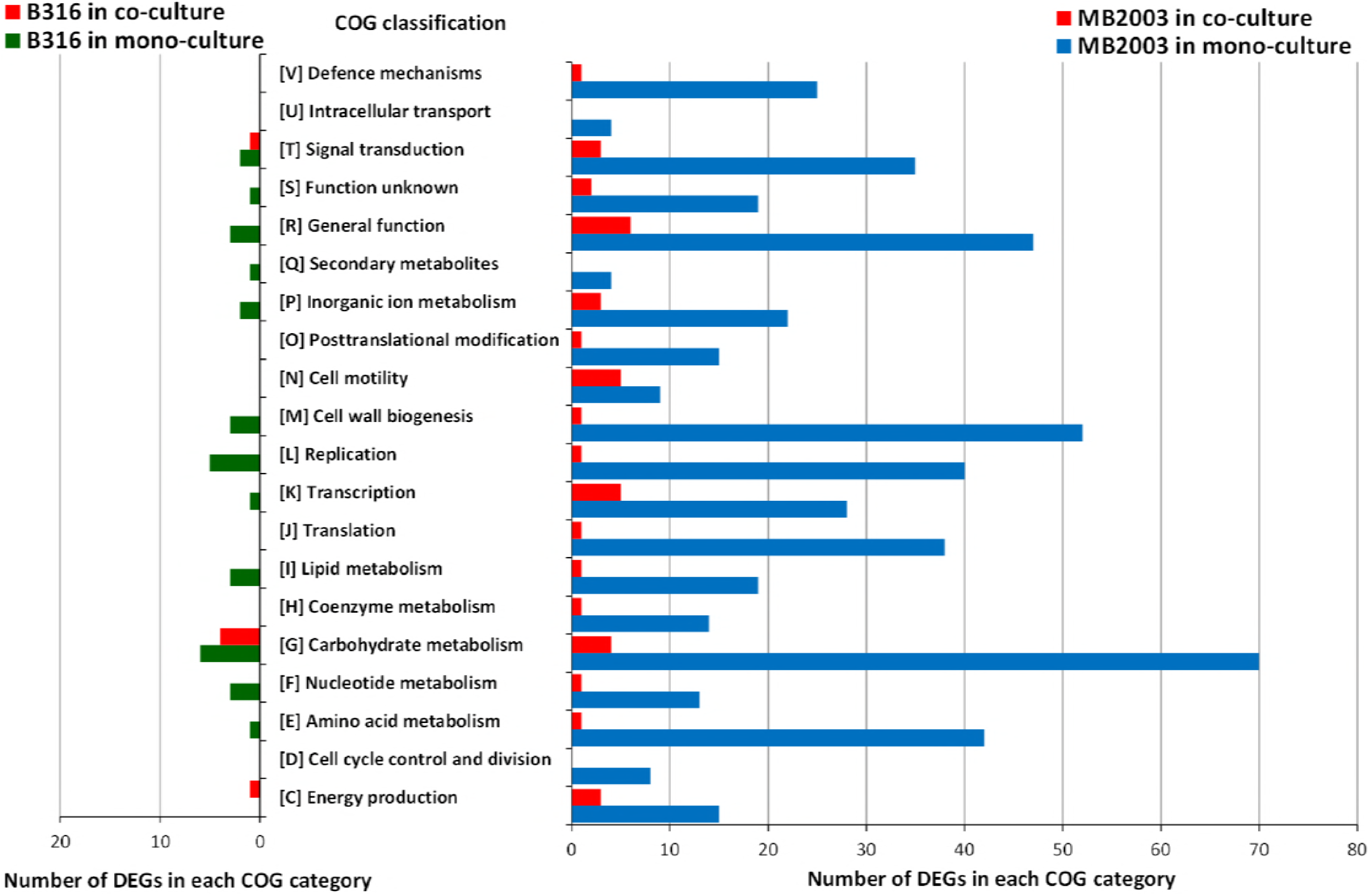
COG classifications of DEGs from *B. proteoclasticus* B316^T^ and *B. hungatei* MB2003 grown in mono- and co-culture on pectin. Analysis includes all DEGs with COG classifications.

### Differentially expressed genes for CAZymes predicted to be secreted to initiate xylan and pectin degradation

RNA sequencing (RNA-Seq) transcriptional analysis of CAZyme assignments for DEGs indicated differences in carbohydrate metabolism of mono- and co-cultures of B316^T^ and MB2003 grown on xylan (predominantly GAX) or pectin (predominantly XGA and RG-I) (Data Sets S3 to S6). The presence of a predicted signal peptide signal sequence on the CAZyme genes indicated that they were membrane bound. Strikingly, for B316^T^ only *xsa43A* (xylosidase/arabinofuranosidase) from xylan-containing mono-cultures was highly up-regulated at 3.36 log_2_-fold change (Fig. 4A). The observed up-regulation of *xsa43A* (bpr_I0302) in B316^T^ mono-cultures grown on xylan (Fig. 4A), signifies that this is an important enzyme for xylan degradation. Previous functional characterization of the *xsa43E* from B316^T^ found this enzyme has dual β-xylosidase and α-L-arabinofuranosidase activities (13), and also encodes an N-terminal GH43 (Pfam04616) catalytic domain, and a C-terminal CBM6 (Pfam03422) non-catalytic module that has been shown to bind xylan in other organisms (14). CBM6 domains are able to recognize xylose either as a monosaccharide, at the non-reducing end of xylo-oligosaccharides, or within the side chain components of xyloglucan (15). Several CBM6 modules also recognize (1,3)-β-D-linkages at the non-reducing end of β-glucans (16, 17), and appear to have co-evolved with their associated catalytic domains to acquire the same substrate specificity (18). α-L-arabinofuranosidases cleave arabinose side chains from substituted xylo-oligosacccharides derived from xylan (19). It is hypothesized that *xsa43A* is secreted into the extracellular environment and has a role in disrupting the complex inter- and intra-polymer networks within the plant cell wall.

**Figure. 4.**
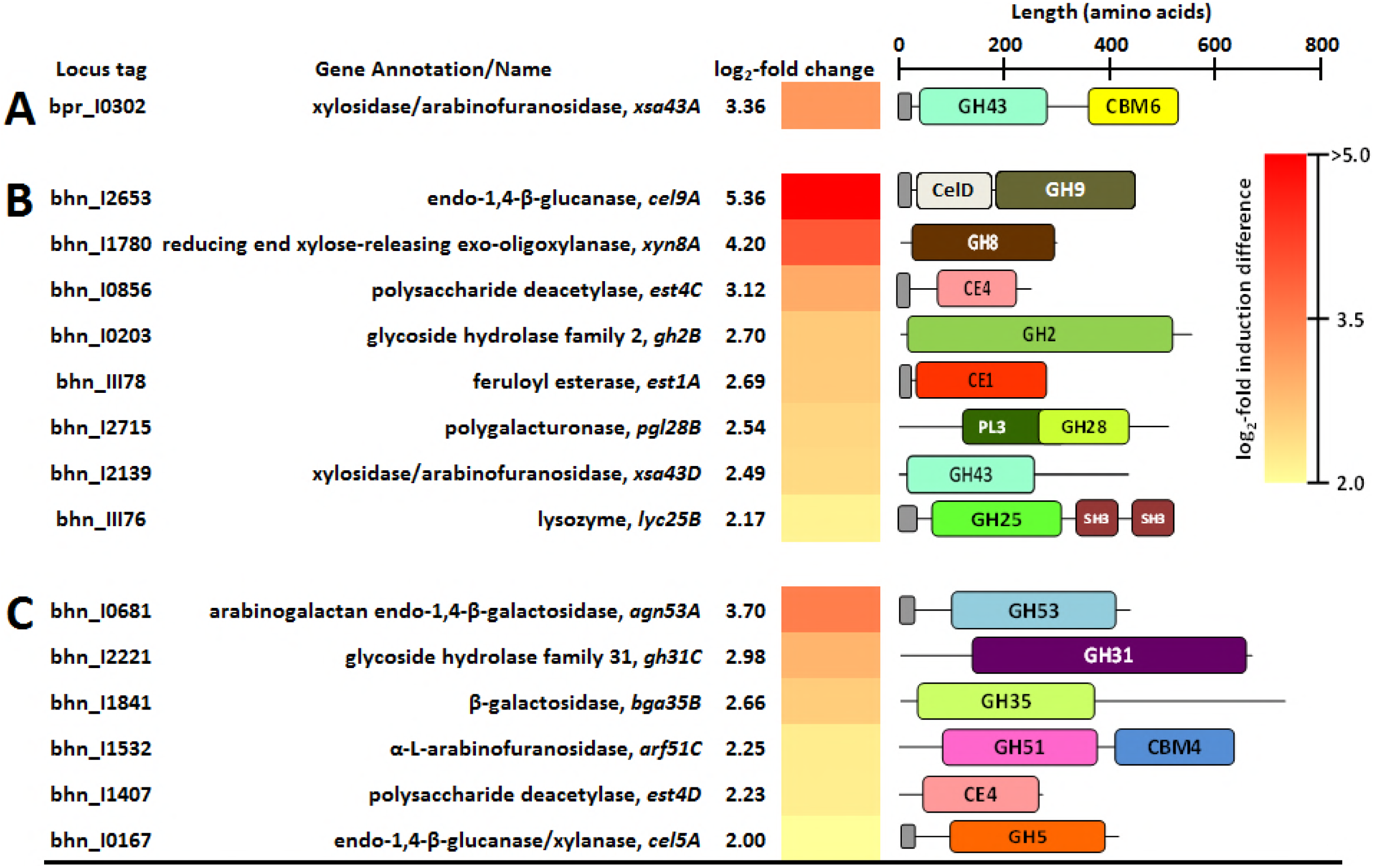
CAZyme encoding DEGs up-regulated in xylan-grown cultures. A, B316^T^ grown in mono-culture; B, MB2003 in mono-culture; C, MB2003 grown in co-culture., signal peptide sequences.

Furthermore, high arabinofuranosidase activity has previously been observed with *B. fibrisolvens* grown on xylan suggesting arabinofuranosidase activity may be regulated by the availability of arabinose (20-23). It was surprising that B316^T^ did not significantly up-regulate any genes encoding secreted CAZymes when grown in mono- or co-culture on pectin, however it displayed better growth in mono-culture on pectin than on xylan (Fig. 1B).

In comparison, MB2003 expressed 6 CAZyme DEGs predicted to be presented on the surface in the xylan-containing cultures and 8 in the pectin-containing cultures (Fig. 4B, C and 5B, C). MB2003 xylan transcriptomes significantly expressed secreted enzymes containing CE1 and CE4, as well as GH53, GH10 and GH5 CAZy domain containing genes (Fig. 4B and C). The extracellular feruloyl esterase *est1A* (bhn_III78) and polysaccharide deacetylase *est4C* (bhn_I0856) predicted to target the acetyl side groups of xylan, were significantly up-regulated in MB2003 xylan mono-cultures (Fig. 4B and Data Set S1). Furthermore, in MB2003 xylan mono-cultures, *cel9A* (endo-1, 4-β-glucanase, GH9/Pfam00759) was the most up-regulated CAZyme gene with 5.36 log_2_-fold change (Fig. 4B). In addition to the secreted enzymes containing the various GH and CE CAZy domains for degradation of GAX and its constituent side groups, the GH13 and PL1 domains that target (1,4)-β-D-linkages within the galacturonic acid backbones are also necessary for the extracellular degradation of XGA and RG-I.

The 8 genes encoding secreted CAZymes involved in pectin breakdown up-regulated in MB2003 mono-culture transcriptomes varied greatly in their predicted enzymatic capability (Fig. 5B). MB2003 significantly expressed secreted enzymes containing GHs 53, 30, 13 as well as 43 and 10 CAZy domain containing genes (Fig. 5B). Among these were genes with additional non-catalytic domains including *lyc25B* lysozyme (GH25 and two SH3 domains) and *xsa43A,* xylosidase/arabinofuranosidase (GH43 and CBM6 domains). The *xsa43A* gene of MB2003 is homologous to the *xsa43A* of B316^T^ (Fig. 4A), and is proposed to possess dual β-xylosidase and α-L-arabinofuranosidase activities. It is hypothesized that *xsa43A* is secreted where extracellular debranching of pectin results in release of arabinose and xylose side groups prior to pectic-oligosaccharide assimilation.

**Figure. 5.**
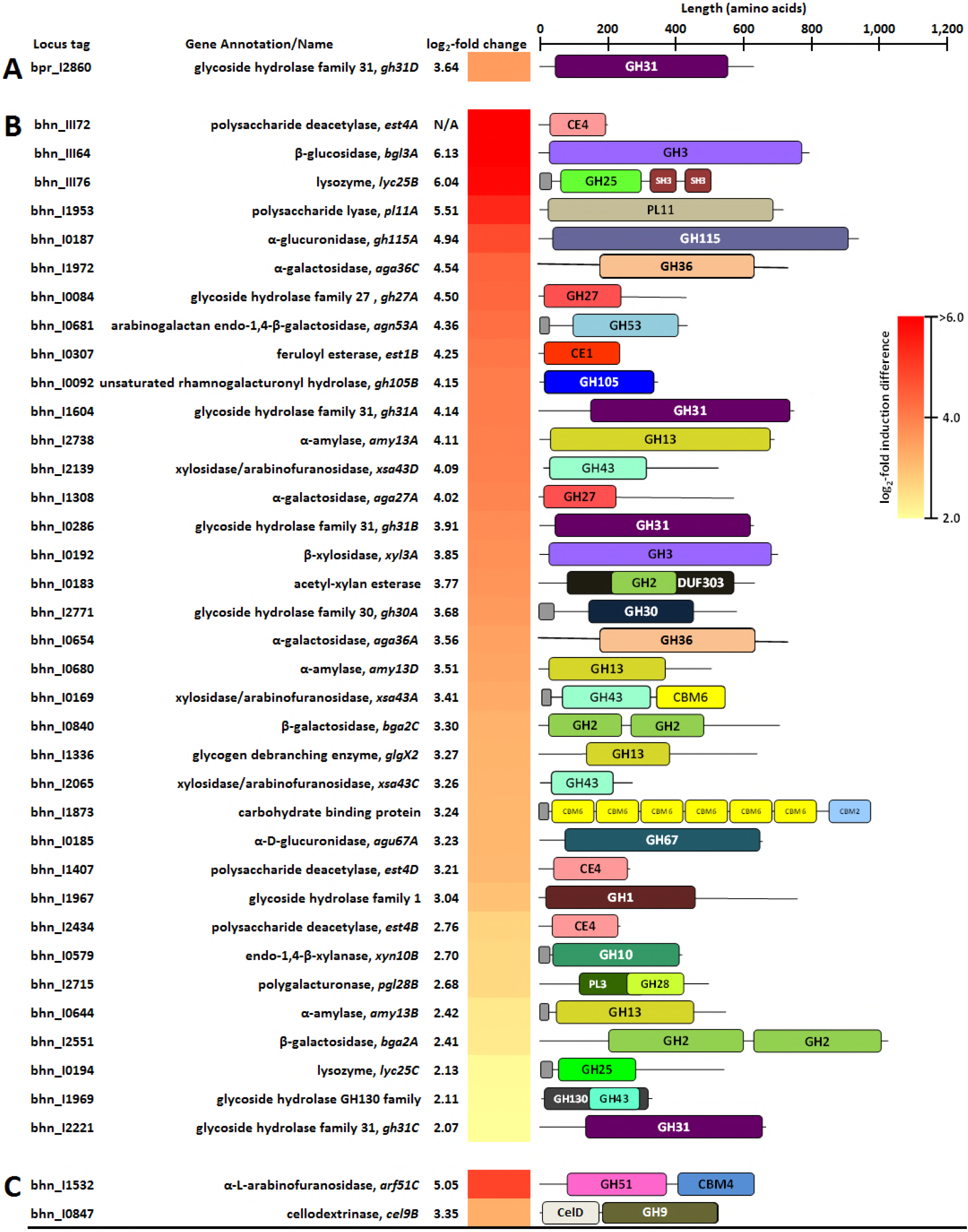
CAZyme encoding DEGs up-regulated in pectin-grown cultures. A, B316^T^ grown in co-culture; B, MB2003 in mono-culture; C, MB2003 grown in co-culture., signal peptide sequences.

The *gh30A* (bhn_I2771) was significantly up-regulated only in MB2003 mono-cultures grown on pectin (Fig. 5B). This enzyme has a GH30 (Pfam02055) catalytic domain that is predicted to have activities in debranching of pectin to release *D*-xylose and to hydrolyse (1,4)-β-D-linkages in xylans (24). However, the functional role of GH30 enzymes, and their contribution to xylan and pectin degradation, will require further biochemical investigation. The β-1,4-galactanases containing GH53 catalytic domains are predicted to degrade galactan and arabinogalactan side chains in the hairy regions of pectin, by attacking the 1,4-β-D-galactosidic linkages in Type I arabinogalactans (25). In MB2003 *agn53A* (bhn_I0681) was significantly expressed in mono-culture pectin-grown cells and was the most up-regulated gene in co-culture xylan-grown MB2003 cells (Fig. 5B and 4C). This suggests that *agn53A* encodes an important xylan- and pectin-degrading enzyme in MB2003. MB2003 encodes one secreted CBP (bhn_I1873, 984 aa), and this protein was significantly up-regulated only in pectin-grown mono-culture cells (Fig. 5B). The domain structure of MB2003 bhn_I1873 is unique, containing six CBM6 domains towards the N-terminus and a single C-terminal CBM2a domain. We propose that the significantly up-regulated and above mentioned enzymes contribute to the surface trimming and depolymerization of complex xylans (predominantly GAX) or pectin (predominantly XGA and RG-I).

### Preferential upregulation of genes encoding surface binding proteins that facilitate uptake of monosaccharides, xylo- and pectic- oligosaccharides via ABC transport systems in response to growth on xylan and pectin

Functional annotations of DEGs assigned to the “Carbohydrate transport and metabolism” COG category [G] were investigated in order to determine which carbohydrates are taken up by B316^T^ and MB2003 cultures grown on xylan and pectin. The transcriptional analysis revealed a substantial number of ABC transporters up-regulated in MB2003 grown on xylan or pectin in both co- and mono-cultures. DEGs associated with ABC system carbohydrate transport and their organization in relation to the surrounding genes encoding polysaccharide-degrading enzymes in the xylan and pectin transcriptomes of B316^T^ and MB2003, are shown in Fig. 7. Strikingly, the most up-regulated genes in both the xylan and pectin transcriptomes encode solute-binding proteins (SBPs) and permease proteins (PPs) of the sugar ABC transport systems that contained a carbohydrate uptake transporter type 1 (CUT1) or a CUT2 domain. The CUT1 domains were represented by COG1653 (SBPs), and COG0395, COG1175 and COG4209 (PPs). The CUT2 domains were represented by COG1879 and COG4213 (SBPs), as well as COG1175 and COG4214 (PPs). The most prevalent belonging to the Carbohydrate Uptake Transporter 1 (CUT1) family that mediate di- and oligo-saccharide uptake (26). The CUT1 and CUT2 family SBPs are represented by the COG1653 and COG4213 domains (27).

**Figure. 6.**
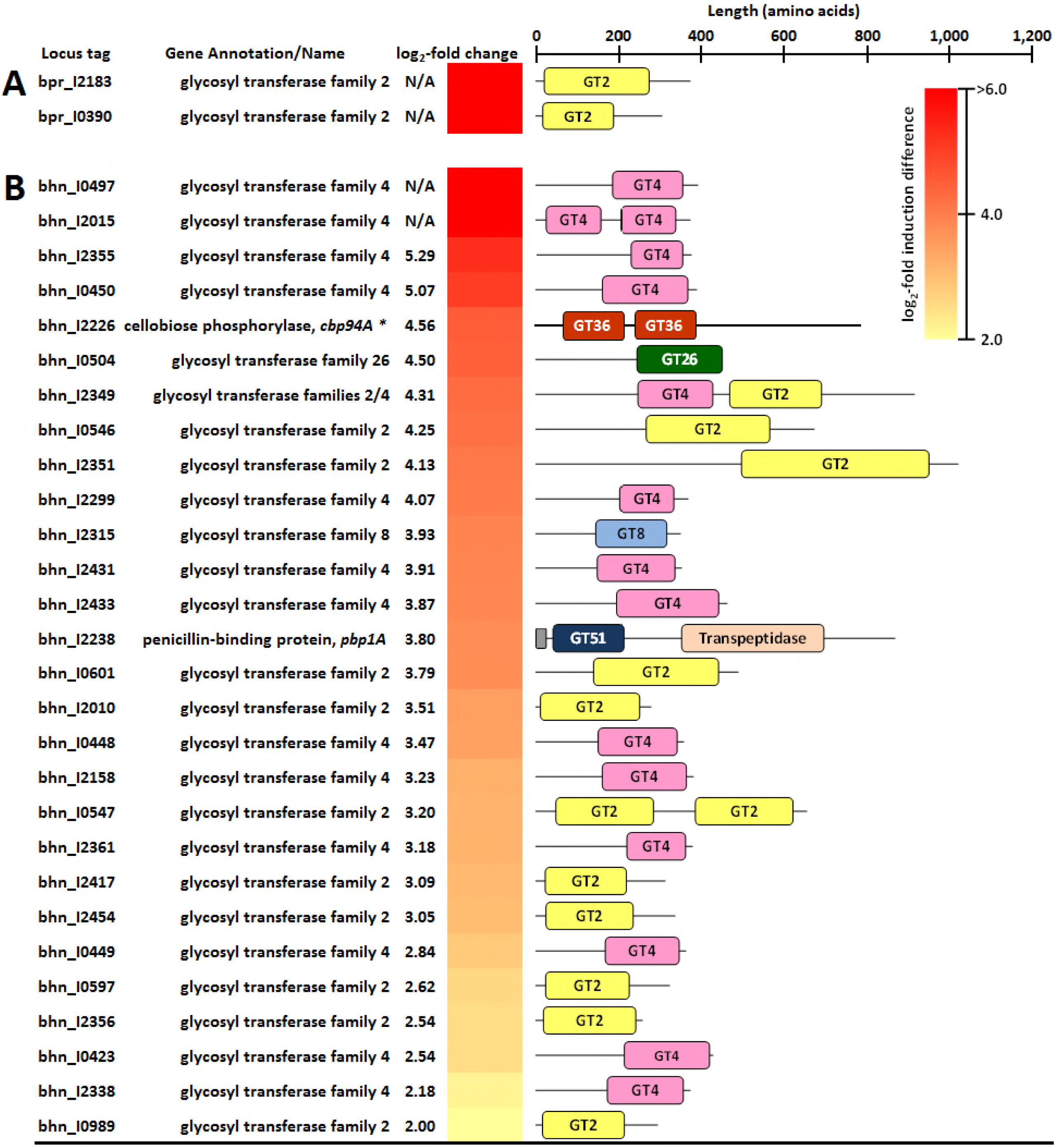
GTs up-regulated during mono-culture growth on pectin. A, B316^T^ grown in mono-culture; B, MB2003 in mono-culture. *, Domains for cellobiose phosphorylase were initially designated as GT36, these have recently been reclassified as GH94 hence the designation in the *cbp94A* gene name., signal peptide sequences.

**Figure. 7.**
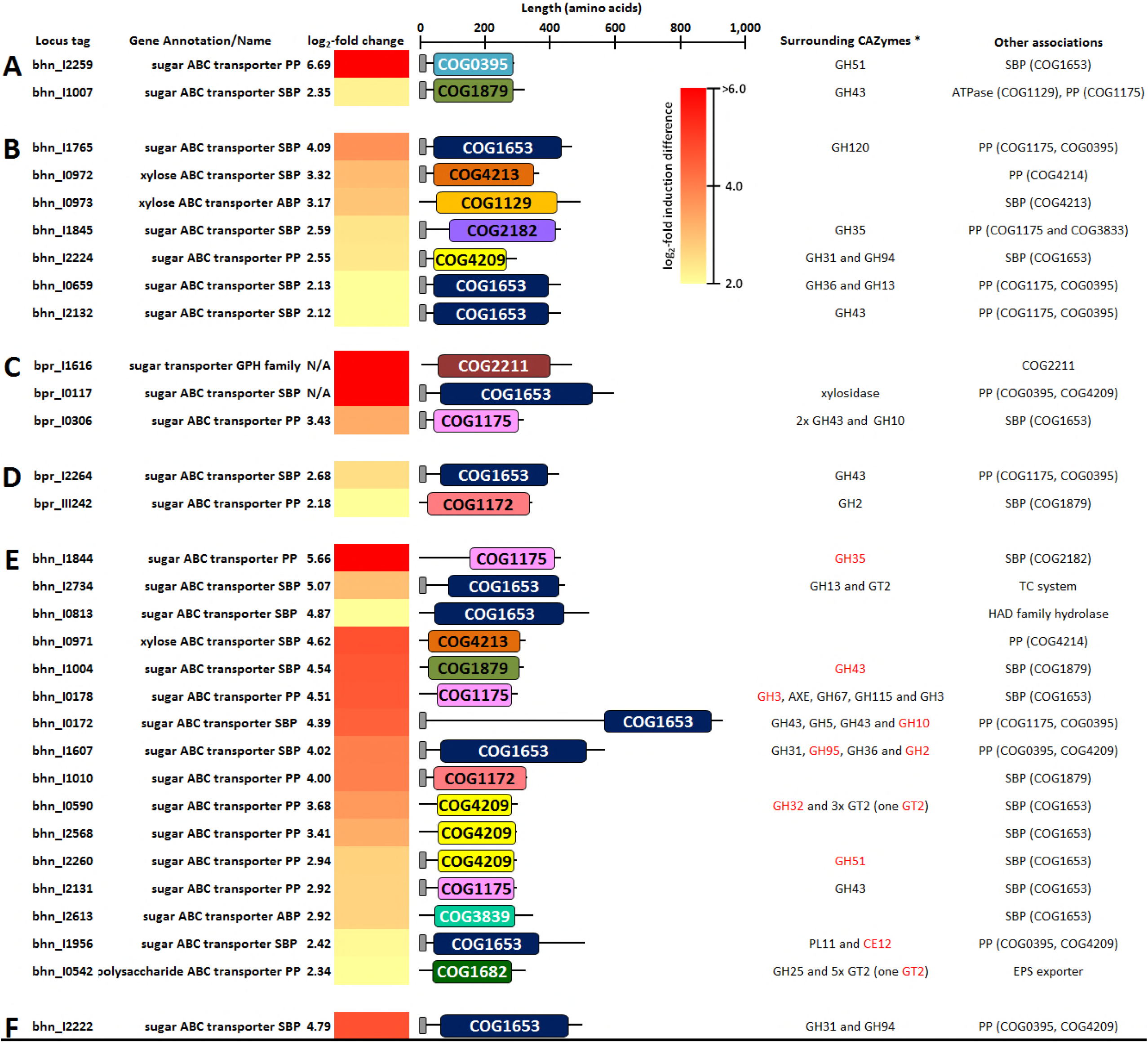
Functional domains of DEGs encoding carbohydrate transport proteins and surrounding CAZymes identified in xylan- or pectin-grown cultures. A, MB2003 mono-culture on xylan; B, MB2003 co-culture on xylan; C, B316^T^ mono-culture on pectin; D, B316^T^ co-culture on pectin; E, MB2003 mono-culture on pectin; F, MB2003 co-culture on pectin. *, CAZyme families of genes and DEGs (red) encoding polysaccharide-degrading enzymes co-localized with carbohydrate transport genes. Abbreviations: SBP, solute-binding proteins; PP, permease protein; AXE, acetyl-xylan esterase; ABP, ATP-binding protein; EPS, exopolysaccharide; TC system, two component system histidine kinase. COG designations: COG0395, COG1175, COG1653 and COG1879, periplasmic component of ABC-type sugar transport system; COG1682, ABC-type polysaccharide/polyol phosphate export systems; COG2182, maltose-binding periplasmic proteins/domains; COG3833, permease component of ABC-type maltose transport systems; COG2211, Na^+^/melibiose symporter and related transporters; COG4213, periplasmic component of ABC-type xylose transport system; COG1129 and COG3839, ATPase component of ABC-type sugar transport systems; COG4209, permease component of ABC-type polysaccharide transport system; COG1172, permease component of ribose/xylose/arabinose/galactoside ABC-type transport systems., signal peptide sequences.

Overall, the transcriptomic analysis identified up-regulation of 45 and 38 genes when grown on xylan, with 10 and 78 genes when grown on pectin, encoding membrane proteins (predicted to function as carbohydrate transporters) in B316^T^ and MB2003 respectively (Data Set S1 and S2). Among the 38 and 78 ABC transporter proteins expressed in MB2003, significant up-regulation of three and four genes encoding proteins with predicted functions as xylose ABC transporters were detected in xylan and pectin transcriptomes respectively. In addition, MB2003 also displayed significant expression of 13 and 46 genes associated with many different transport systems and target substrates in the xylan and pectin transcriptomes, respectively. For the B316^T^ transcriptomes, no genes encoding xylose ABC transporter systems were differentially expressed and the only genes with predicted functions as sugar ABC transporters were found.

While numerous DEGs (log_2_ fold change >2) associated with ABC system carbohydrate transport were found in MB2003 xylan or pectin transcriptomes (Fig. 7A, B, E and F), SBPs and PPs were identified in the B316^T^ pectin transcriptomes only. In total, five DEGs with predicted functions as sugar ABC transporters were found for B316^T^ grown on pectin; three genes in the mono-culture, and two genes in the co-culture condition (Fig. 7C and D). Signal peptides were predicted for the majority of sugar ABC transporter SBPs and PPs identified to be significantly upregulated, and in MB2003 for the xylose transporter SBPs. Detection of a large number of significantly, differentially expressed ABC transporter SBP-encoding genes in B316^T^ and MB2003 signifies the importance of SBP-dependent ATP-driven active transport of oligo- and mono-saccharides for growth. The preference of SBPs associated with oligo- and monosaccharide-specific ABC transporters correlates with the concentrations of xylose and galactose in B316^T^ and MB2003 co-cultures detected using IC (Fig. 1), suggesting an efficient uptake of oligomers produced by the initial breakdown of complex xylan and pectin.

### Co-localization of CAZymes with highly expressed ABC transporter binding proteins reveals interplay between polysaccharide utilization machinery

The polysaccharide-degrading enzymes found adjacent to these SBPs include a number of xylosidases/arabinofuranosidases (GH43), endo-1,4-β-xylanase (GH10), and a β-mannosidase (GH2). MB2003 cells grown on xylan and pectin up-regulated 9 and 17 genes respectively, associated with sugar ABC transport systems and target substrates (Fig. 7). Examples of the surrounding polysaccharide-degrading enzymes found in MB2003 xylan grown cells include: α-L-arabinofuranosidase (GH51), xylosidases/arabinofuranosidases (GH120, GH30 and GH43), α- and β-galactosidases (GH35 and GH36), sucrose phosphorylase (GH13), and glycosidases (GH31) (Fig. 7A and B). MB2003 grown in co-culture on xylan and mono-culture on pectin up-regulated bhn_I0972 and bhn_I0973, and bhn_I0971 genes respectively, associated with xylose transport (Fig. 7B and E). All of the SBPs represented by the COG1653 domain in both B316^T^ and MB2003 were found co-localized with other genes predicted to be involved in xylan degradation and metabolism, including various genes encoding GH43, GH10, GH51, GH53 and CE12 CAZy domains (Fig. 7A and B). This includes two SBP-encoding genes identified in xylan-grown B316^T^ mono- and co-cultures, bpr_I0182 and bpr_I0313, which were previously reported in B316^T^ xylan-grown cells as being part of a polysaccharide utilization loci or PULs (8), predicted to be involved in hemicellulose degradation. The PULs representative of these SBPs contain genes encoding β-xylosidases, α-glucuronidases, acetyl-xylan esterases, ferulic acid esterases and secreted endo-1, 4-β-xylanases (24), and are thought to be important for efficient hemicellulose metabolism by B316^T^.

A variety of polysaccharide-degrading enzymes have been identified proximal to the genes up-regulated in MB2003 pectin transcriptomes and examples include: xylosidases/ arabinofuranosidases (GH43, GH51), endo-1,4-β-xylanase (GH10), acetyl-xylan esterase, endo-1,4-β-glucanase/xylanase (GH5), β-glucosidase (GH31), α-D-glucuronidase (GH67), α-glucuronidase (GH115), β-xylosidase (GH30), pectin eterase (CE12) and rhamnogalacturonan lyases (PL11) (Fig. 7E). Also, a total of nine GT2 genes were also identified proximal to the SBPs (Fig. 6B). The CAZymes surrounding the various SBPs were highly up-regulated and many of these were only expressed in the mono-culture MB2003 transcriptomes (Data Set S3). In contrast, only one gene (bhn_I2222) was significantly up-regulated in co-culture, in proximity to a glucosidase (GH31) and cellobiose phosphorylase (GH94) (Fig. 7F). Overall, SBPs and PPs with the COG1653 were the most abundant domains identified in MB2003 pectin grown cells. These findings are consistent with the extracellular breakdown of xylan and pectin followed by capture and uptake of monosaccharides, xylo- and pectic- oligosaccharides by the ABC transporter.

### Cytosolic enzymatic debranching of xylo- and pectic- oligosaccharides

The polysaccharide degrading machinery of both MB2003 and B316^T^ appear to be optimized to maximize intracellular breakdown, leading us to hypothesize that *Butyrivibrio* may adopt a ‘selfish’ strategy for hemicellulose and pectin breakdown in the rumen. The xylan- and pectin-derived xylo- and pectic- oligosaccharides entering the cytoplasm are degraded in concert by a large suite of exo-acting enzymes. The highly expressed genes predicted to encode oligosaccharide debranching enzymes (and their catalytic domains) are: *xyl120A, B* (GH120), *est1B, D, E* (CE1), *xsa43B-I* (GH43), *xyn8A* (GH8), *xyl3A* (GH3), *agu67A* (GH67), *arf51A-C* (GH51), *gh115A* (GH115), *gh31B, C* (GH31), *est2A* (CE2), *est4A-E* (CE4), and an unclassified acetyl-xylan esterase (AXE) (Figure 6.2C). The strong up-regulation of debranching enzymes and sugar SBP-dependent ABC transporter systems, indicates that extracellular degradation of xylan does not result in complete removal of side chain groups and that the xylo-oligosaccharides are assimilated into the cytoplasm for further degradation by intracellular enzymes.

Surprisingly, B316^T^ significantly up-regulated only *gh31D* (a possible α-glucosidase (EC 3.2.1.20) or α-xylosidase (EC 3.2.1.-)) with a log_2_-fold change of 3.64, when grown in co-cultures on pectin (Fig. 5A). The two most up-regulated cytosolic CAZyme genes in MB2003 xylan transcriptomes were *xyn8A* (reducing end xylose-releasing exo-oligoxylanase, GH8/Pfam01270) and *est4C* (polysaccharide deacetylase; CE4/Pfam01522), each with > 3.0 log_2_-fold change (Fig. 4B). The 36 CAZyme genes up-regulated in MB2003 pectin-grown mono-culture varied greatly in their predicted enzymatic capability (Fig. 5B) and included genes with multiple domains (acetyl-xylan esterase containing a GH2 sugar-binding domain and domain of unknown function DUF303 (Pfam03629)), genes containing two identical catalytic domains within the same gene, (β-glucosidase *bgl3D* with two GH3 domains (Pfam00933)) and the two β-galactosidases *bga2A* and *bga2C* which each contain two GH2 (Pfam00703) domains. The polysaccharide lyase, *pl11A* (PL11/COG14111 domains), also known as rhamnogalacturonan lyase (EC 4.2.2.-) or exo-unsaturated rhamnogalacturonan lyase (EC 4.2.2.-), was highly up-regulated in the MB2003 pectin-grown mono-culture with a log_2_-fold change of 5.51.

Examples of genes with additional non-catalytic domains include *lyc25B* lysozyme (GH25 and two SH3 domains) and *xsa43A,* xylosidase/arabinofuranosidase (GH43 and CBM6 domains). DEGs encoding GT CAZy functions were only found in the pectin-grown mono-cultures for both B316^T^ and MB2003 (Fig. 6). Two GT domain-containing genes were found for B316^T^ (Fig. 6A), compared to 28 in MB2003 (Fig. 6B). The two GT genes found in B316^T^ both possess a single GT2 (Pfam00535) domain, approximately 400 aa in length. The GTs of MB2003 consist of ten GT2, thirteen GT4, and single genes containing GTs 8, 26, 51, 36 and both GT2 and GT4 domains. To summarize, using the expression and biochemical data presented in this work, Fig. 8 shows the proposed models for the primary degradation, transport and cytosolic breakdown by *B. proteoclasticus* B316^T^ and *B. hungatei* MB2003 of the different classes of xylan (GAX) and pectin (XGA and RG-I) presented to the rumen microbiota.

**Figure. 8.**
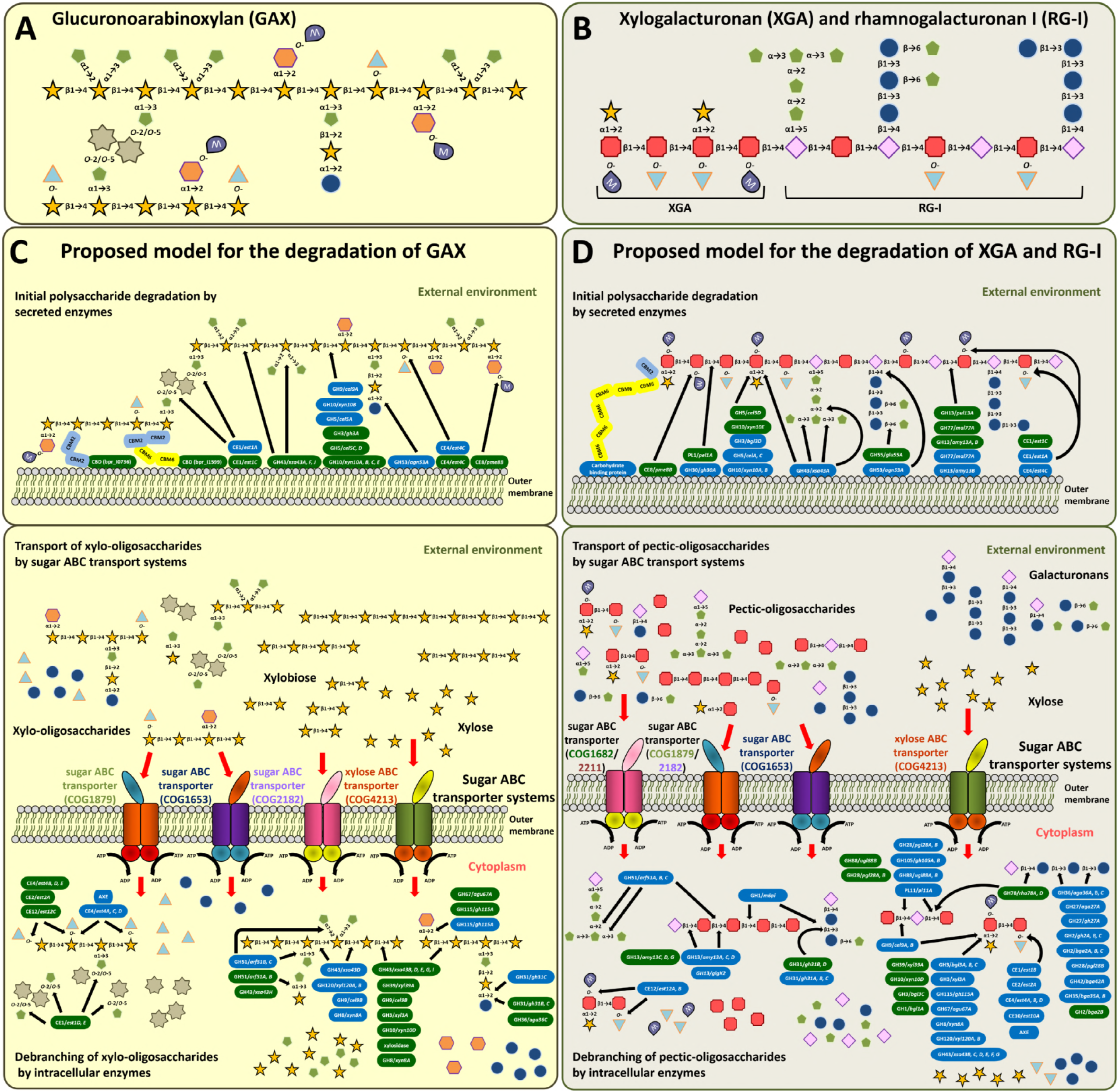
Proposed models for the degradation of different forms of xylan and pectin by *B. proteoclasticus* B316^T^ and *B. hungatei* MB2003. In the upper panels the monosaccharides, side groups, and linkages in the main classes of A, xylan (GAX) and B, pectin (XGA and RG-I), are represented. Schematic diagrams of the structures of the main classes of xylan and pectin. The A, xylan (GAX) and B, pectin (XGA and RG-I) structures shown are not quantitatively accurate. The proposed models for degradation of complex GAX (C) along with XGA and RG-I (D) includes the primary attack, assimilation and intracellular processing. The black arrows indicate examples of the linkages cleaved by the enzymes. The significantly up-regulated polysaccharidases are represented by ovals shown in green for B316^T^ and blue for MB2003. Enzymes are identified based on their CAZy classifications, consisting of GHs, CEs, PLs and CBMs. The red arrow heads indicate xylo- or pectic-oligosaccharide transport between cellular locations. SBPs signify the surface sugar and xylose binding proteins of the ABC transporter systems. Schematic representation of the ATP-driven sugar ABC transport systems likely to mediate the uptake of xylan- or pectin-derived soluble sugars are also included. All polysaccharidases and transport systems shown are based on transcriptional analysis of genes deemed significantly expressed in either mono- or co-culture growth conditions. Symbols: 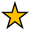, *D*-xylose; 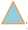, acetyl group; 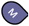, methyl group;, *L*-arabinose; 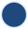, *D*-galactose; 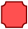, *D*-galacturonic acid; 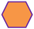, *D*-glucuronic acid; 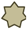, ferulic acid; 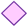, *L*-rhamnose. Abbreviations: AXE, acetyl-xylan esterase; CBD, carbohydrate binding protein. COG designations: COG1653, ABC-type sugar transport system; COG1682, ABC-type polysaccharide/polyol phosphate export systems; COG1879, ABC-type sugar transport system; COG2182, maltose-binding periplasmic proteins/domains; COG2211, Na^+^/melibiose symporter and related transporters; COG4213, ABC-type xylose transport system. The pectin structures shown are not quantitatively accurate.

## DISCUSSION

Xylan and pectin are the most abundant plant structural polysaccharides after cellulose, and are major sources of energy for microbial fermentation within ruminants. The xylan derived from oat-spelts used in this experiment, is largely composed of GAX which is comprised of a xylose monomer backbone (28). The pectin used for this experiment was derived from apple and is largely comprised of RG-I and XGA, with backbones composed of galacturonic acid and rhamnose monomers (29). Rumen bacteria degrade xylans to xylose and arabinose and pectins are degraded to predominantly, rhamnose and galactose, and a variety of higher oligosaccharides are produced from both substrates (30-32). The qPCR analysis of mono-culture samples confirmed *B. proteoclasticus* B316^T^ as a strong degrader of xylan and pectin, while *B. hungatei* MB2003 was unable to utilize either substrate to support growth (Fig. 1). MB2003 was only able to grow in co-cultures with B316^T^, utilizing the sugars released by B316^T^ from xylan or pectin. The relationship between MB2003 and B316^T^ is not simple commensalism, as the growth of B316^T^ cells in co-cultures with MB2003 was compromised compared to their mono-culture growth. The relationship appears to be a kind of competitive parasitism in which B316^T^ acts as a primary degrader providing soluble sugars, allowing MB2003 to compete for the soluble sugars in a parasitic manner.

Analyses of the soluble sugars released during growth showed that fermentation in the B316^T^ mono- and co-culture samples was complete by 12 h, after which the levels of xylose decreased dramatically (Fig. 1A). The complete absence of released monosaccharides in MB2003 samples supports the VFA and pH experimental evidence for its inability to utilize xylan in mono-culture (Fig. 1A). The total pool of xylose detected in the B316^T^ + MB2003 co-cultures on xylan and the reduced xylose concentration at 12 h post-inoculation, was not as dramatic as in the B316^T^ mono-culture (Fig. 1A), apparently due to the presence of MB2003 cross-feeding on the released xylose. It is possible that the other sugars analyzed (or not detected) could have been released and used immediately (or at time points not analyzed) such that their concentrations in the samples did not appear to increase. However, this seems unlikely given that xylan extracted from oatspelts is typically >70% xylose and <10% arabinose and <15% glucose (CAS Number 9014-63-5, X0627, Sigma-Aldrich).

In pectin-grown mono- and co-cultures, galactose was the predominant monosaccharide detected. B316^T^ grown in mono-culture contained the highest concentrations (Fig. 1B) which is consistent with B316^T^ being more efficient at breaking down pectin. This is in contrast to the very poor, if any, ability of MB2003 to utilize this substrate (11). However, the rate of galactose utilization after the early log phase was similar between the mono-cultures and co-culture, suggesting that MB2003 could compete with B316^T^ for the released galactose. Also, it is likely that galacturonic acid, not detected in the IC analysis, was released during hydrolysis of pectin and may have been used by B316^T^, while not by MB2003. In a previous study on B316^T^ grown on pectin (33), an increase in, rhamnose, arabinose and other mono- and disaccharides was observed, but not galacturonic acid or galactose. The galacturonic acid and galactose released from hydrolysis of pectin in the inoculum may have been consumed rapidly and hence not detected.

The growth experiments showed that MB2003 entered a state of ‘starvation’ as it was unable to utilize xylan or pectin when grown in mono-culture, but was capable of significant growth on both substrates when co-cultured with B316^T^. MB2003 appears to sustain itself in co-culture until primary degradation of xylan and pectin by B316^T^ is underway, allowing for the uptake of released soluble carbohydrates. The transcriptome analysis showed that in B316^T^, only genes required for substrate utilization were significantly up-regulated (Fig. 2 and 3). Genome sequence information (8) and proteome analyses (34, 35) indicate that B316^T^ primarily attacks the xylan backbone and main substituent groups of hemicellulose outside the cell via secreted enzymes. The variable length substituted or un-substituted xylo-oligosaccharides are then thought to be transported into the cell where the final degradation occurs. B316^T^ applies a similar approach for the degradation of pectin, but the pectin backbone is composed of galacturonic acid. The comprehensive enzymatic machinery of these rumen bacteria allow for the highly efficient utilization of internalized xylo- and pectic-oligomers in the cytoplasm, without loss to competing rumen species.

*B. hungatei* MB2003 does not initiate the breakdown of the backbones of either xylan or pectin, but appears to act as a competitor for sugars released from the insoluble substrates through the primary degradation activity of B316^T^. MB2003 mono-cultures in xylan or pectin-containing media, appear to enter a state of ‘starvation’ as there was no significant growth, but its cells strongly up-regulated genes involved in almost every biological process to scavenge substrates in an attempt to initiate growth. MB2003 grown in co-culture with B316^T^ on xylan or pectin also shows up-regulation of many genes, but to a lesser extent than in the monocultures. These upregulated genes include those encoding the enzymatic machinery required to utilize the oligosaccharides released from the initial degradation of xylan and pectin by B316^T^. The initial degradation of xylans and pectins by B316^T^ will also effect the rumen microbial ecosystem by promoting the growth of secondary degraders, *Butyrivibrio* and *Pseudobutyrivibrio*, which will lead to butyrate production, a key animal health promoting fermentation end-product.

The B316^T^ and MB2003 gene compliments suggest they cannot completely degrade complex GAX, XGA and RG-I to monosaccharides on the outside of their cells, and implies that they must transport a variety of substituted xylan and pectin oligomers across their bacterial cell walls. The degradation of these oligomers into their constituent monomers is achieved through the activity of several classes of cytosolic enzymes, including β-xylosidases, α-galactosidases and α-glucuronidases that contain GH3, GH27, GH115 and GH67 CAZy domains. The starved MB2003 cells inoculated in mono-cultures on pectin, significantly expressed intracellular genes encoding enzymes containing GH3, GH27, GH115 and GH67 CAZy domains (Fig. 5B).

In order to make use of oligo- and monosaccharides derived from extracellular digestion of lignocellulosic material, it is necessary for the released soluble sugars to be transported into the cell. *Butyrivibrio* have Gram-positive cell wall structures (10-12), and monosaccharide transport across the bacterial cell wall is mediated by a variety of extracellular SBPs linked to dedicated sugar ABC transporter systems. Recent data on the B316^T^ secretome (34, 36) and carbohydrate transport-associated membrane proteins (35), as well as genome sequence analysis of MB2003 (10, 11), identified a large number of sugar-specific ABC transporter SBPs.

MB2003 was able to compete with B316^T^ for both xylan and pectin during the log-phase (Fig. 1). Overall, analysis of up-regulated SBPs and their functional domains, along with surrounding genes encoding polysaccharide-degrading enzymes, revealed that the most prevalent functional category was the COG1653 domain that is known to be associated with oligosaccharide transport (35). In B316^T^ and MB2003 pectin transcriptomes, up-regulation of the genes predicted to encode sugar ABC transporter PPs (COG1172) and SBPs (COG1879) respectively, have functional roles as ribose, xylose, arabinose and galactose ABC transporters, suggesting a preference for these substrates. The substantial up-regulation by MB2003 of sugar ABC transport system genes and a large variety of co-localized genes encoding polysaccharide-degrading enzymes (Fig. 7), supports the findings that MB2003 is able to grow only in co-culture on xylan and pectin through cross-feeding on the released oligosaccharides and monosaccharides such as xylose, arabinose and rhamnose (Fig. 8). In addition to the SBPs represented by the COG1653 domain in both B316^T^ and MB2003 found co-localized with other genes involved in xylan degradation and metabolism, genes containing GH13 and PL11 (only in MB2003) CAZy domains (Fig. 7C to F), were expressed in pectin-grown cells. This implies that the availability of a particular carbon source causes activation of a wider network of genes that enable these rumen bacteria to breakdown, transport and metabolize such substrates for growth.

Gene expression in B316^T^ was mostly unaffected by the presence of MB2003, but the transcription level of many CAZymes, including *xyn10D* (bpr_I1083), *xyn10E* (bpr_I1740), and *cel5D* (bpr_I0728) genes that encode endo-1,4-β-xylanase and endo-1,4-β-glucanase activities, respectively, as well as sugar ABC transport system genes, were elevated in B316^T^ (Data Set S1). Xylan polymers are cross-linked via ester linkages with the phenolic acids, ferulic and *p*-coumaric acids (Fig. 8A), which are esterified to the arabinose side chains (37). These linkages have been proposed to account for much of the steric hindrance to plant fibre degradation in the rumen (38), and there is significant interest in rumen microorganisms that possess feruloyl and *p*-coumaroyl esterases that belong to the CE1 (Pfam00756) CAZy family. These esterases function to remove the ferulic acid, methyl and acetyl groups from the xylan polymers and/or oligomers, thus making arabinose and xylose available for growth. The cell-associated β-xylosidases and α-L-arabinofuranosidases are then likely to target the arabinoxylan oligomers from GAX that are subsequently assimilated into the cell. A previous study has shown that high levels of extracellular cinnamoyl esterases are characteristic of a selection of fibre-degrading ruminal bacteria and in particular are common amongst the xylanolytic *B. fibrisolvens* (39). It has been argued that the well-developed xylanolytic and cinnamoyl esterase systems of ruminal fungi give them a distinct advantage over fibrolytic bacteria in the rumen (40). The feruloyl and *p*-coumaroyl esterase activities of *Butyrivibrio* thus warrants further investigation in future work.

Recent studies have shown that in discrete regions of plant cell walls, initial enzymatic attack of pectin increases the access of CBMs to cellulose (41), effectively loosening the polysaccharide interactions to reveal the xylan and xyloglucan substrates (42, 43). This initial stage in enzymatic saccharification of plant cell walls termed amorphogenesis (44), is a possible role of such CBPs containing multiple non-catalytic domains. In the rumen, B316^T^ and MB2003 may thus secrete these non-catalytic CBPs along with polysaccharide-active enzymes as a mechanism to enhance the rate and extent of plant cell wall degradation by disrupting the interface between other polysaccharides. The diversity and variability of CBMs appear to be a signature of extracellular enzymes and non-catalytic CBPs from rumen *Butyrivibrio* (45), and therefore are of interest and future *in vitro* experimentation aimed at investigating the function of CBMs is justified.

The transcriptome analyses have highlighted the differences in the relative amounts of up-regulated genes in *B. proteoclasticus* B316^T^ and *B. hungatei* MB2003 grown in mono- and co-culture that support the observed commensal interactions between the two rumen bacteria in the co-culture growth experiments. These gene expression profiles indicate that B316^T^ grown on either xylan or pectin, showed relatively little change in gene expression when co-cultured with MB2003. In contrast, MB2003 showed poor utilization of either xylan or pectin for growth. These analyses also confirmed the expression of the diverse repertoire of genes that MB2003 and B316^T^ possess which encode polysaccharide-degrading enzymes and sugar ABC transport systems during xylan and pectin degradation. The SBP-mediated ABC-assimilation of polysaccharide-derived soluble sugars, such as xylo- and pectic- oligosaccharides, has been shown to be an essential component for xylan and pectin degradation and utilization by B316^T^ and MB2003. These findings suggest that liberation of GAX, XGA and RG-I from the surrounding polymers also requires numerous non-polysaccharidases, identified in the co-culture experiment. For future work, biochemical characterization of the precise substrate specificities of the SBPs for the various xylo- and pectic- oligosaccharides and characterization of the carbon catabolite repression mechanisms will be essential to defining the sugar transporting systems of B316^T^ and MB2003.

## MATERIALS AND METHODS

*B. hungatei* MB2003 and *B. proteoclasticus* B316T (46) were isolated from the rumen contents of fistulated Friesian dairy cattle as described by S. Noel (47) and sequenced as described by N. Palevich (10, 11) and Kelly *et al*., (8), respectively. Mono- and co-cultures of *Butyrivibrio* strains were grown in media containing 0.5% (v/v) inoculum and supplemented with either 0.5% (w/v) of xylan from oat spelts (Sigma-Aldrich) or pectin isolated from apple (Sigma-Aldrich) as the main carbohydrate source. An overview of the MB2003 and B316^T^ co-culture growth experiment, is presented in Fig. S1. On collection of each sample, 2 mL of each culture was removed and stored at -85 °C for downstream qPCR, HPIC and VFA analyses. The remainder of cultures was snap-frozen and total RNA was extracted using a modified version of a liquid N_2_ and grinding method (48). A comparison of the complete MB2003 and B316^T^ genomes to identify target genes for the real time qPCR assays was conducted using Differential BLAST analysis (DBA) (49). The specificity of each primer/probe assay was demonstrated by qPCR using primers in Table S3. VFA production profiles of *Butyrivibrio* strains grown on insoluble substrates were determined using gas chromatography for the quantification of acetic, butyric and propionic acids and the branched chain acids (BCVFAs) isobutyric and isovaleric acids (50). An additional down-scaled method (51) was used to derivatize formic, lactic and succinic acids. The monosaccharides released during *Butyrivibrio* mono-culture and co-culture growth were determined using a high-pressure ion chromatography (HPIC) method (45). Paired-end libraries of total RNA samples were sequenced with a 200-bp read length on an Illumina HiSeq2000 instrument at the Beijing Genomics Institute (BGI, China). Sequence data was filtered using DynamicTrim (52) and sequence quality was assessed using FastQC (53). Bowtie 2 (54) was used with default parameters, to remove any sequence reads aligning to ribosomal RNA, transfer RNA and non-coding RNA sequences. Reference-based transcriptome assembly (55, 56) was performed because high-quality genome sequences were available for both *B. hungatei* MB2003 (11) and *B. proteoclasticus* B316^T^ (8). Rockhopper (57, 58) was used on the remaining reads to identify differential expression between mono- and co-culture growth of MB2003 and B316^T^ on xylan and pectin conditions separately. An overview of the RNA-seq *in silico* analysis, is presented in Fig. 9. Standard univariate and multivariate statistical tests were performed using R software to analyze the RNA-seq datasets (Table S4, Fig. 9, Data Set S1). Accession numbers for all sequences used in this study and detailed methods are available in the supplemental material (Text S1).

**Figure. 9.**
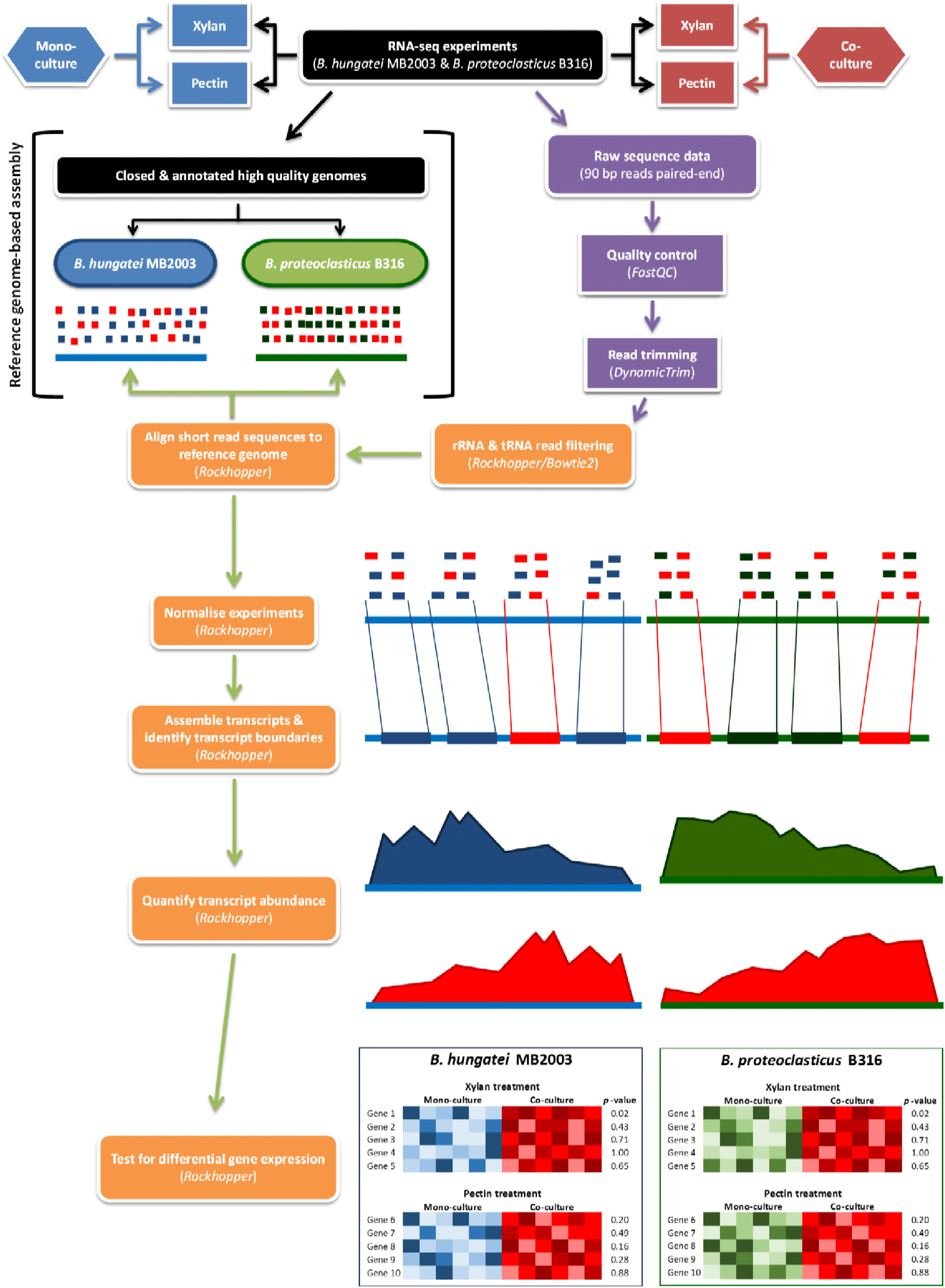
Overview of the workflow for RNA-seq *in silico* analysis. Software and programmes used at each stage are *italicized* and shown in brackets.

## SUPPLEMENTAL MATERIAL

Supplemental material for this article may be found at TBA.

TEXT S1, DOCX file, 0.03 MB.

FIG S1, TIF file, 1.1 MB.

FIG S2, TIF file, 0.6 MB.

TABLE S1, DOCX file, 0.03 MB.

TABLE S2, DOCX file, 0.01 MB.

TABLE S3, DOCX file, 0.01 MB.

TABLE S4, DOCX file, 0.02 MB.

DATA SET S1, XLSX file, 0.04 MB.

DATA SET S2, XLSX file, 0.03 MB.

DATA SET S3, XLSX file, 0.06 MB.

**Accession number(s).** Annotated *B. hungatei* MB2003 and *B. proteoclasticus* B316^T^ genomes were submitted to GenBank under GenBank accession numbers CP017831, CP017830, CP017832, CP017833, and CP001810, CP001811, CP001812, CP001813.

## ACKNOWLEDGEMENTS

We thank Sarah Lewis for assistance with the fermentation end-product analysis, Don Otter for the high-pressure ion chromatography analysis and Eric Altermann for the FGD analysis.

---

**Fig. S1.** Overview of *B. hungatei* MB2003 and *B. proteoclasticus* B316^T^ co-culture growth experiment. * 0.5% (v/v) co-culture inoculum comprised of 0.25% (v/v) B. hungatei MB2003 and 0.25% (v/v) B. proteoclasticus B316 inocula.

**Fig. S2.** Ordinate plots comparing BH-adjusted ANOVA and BH-adjusted KW analyses. A, *B. proteoclasticus* B316^T^ and B, *B. hungatei* MB2003 xylan and pectin complete transcriptome datasets. Abbreviations: BHaov2t *p*-value, Benjamin Hochberg adjusted ANOVA *t*-test *p*-values; BHkrus *p*-value, Benjamin Hochberg adjusted KW analysis of variance by ranks *p*-values.

